## Supplementary figures and images for "Regulation of transcription elongation anticipates alternative gene expression strategies across the cell cycle"

### S1 Fig

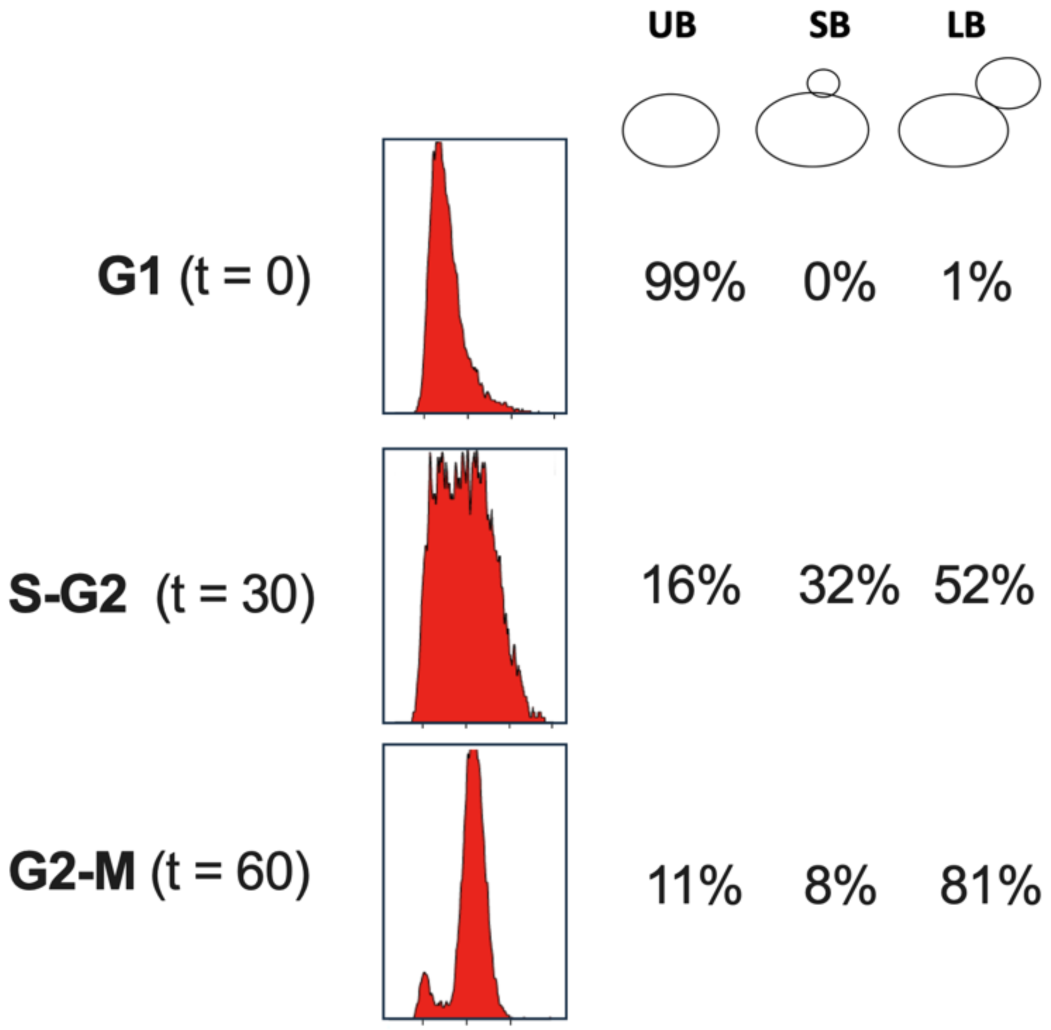

### S2 Fig

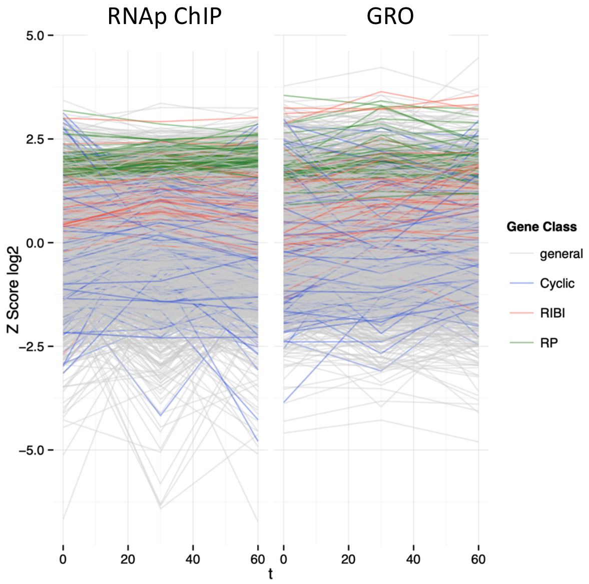

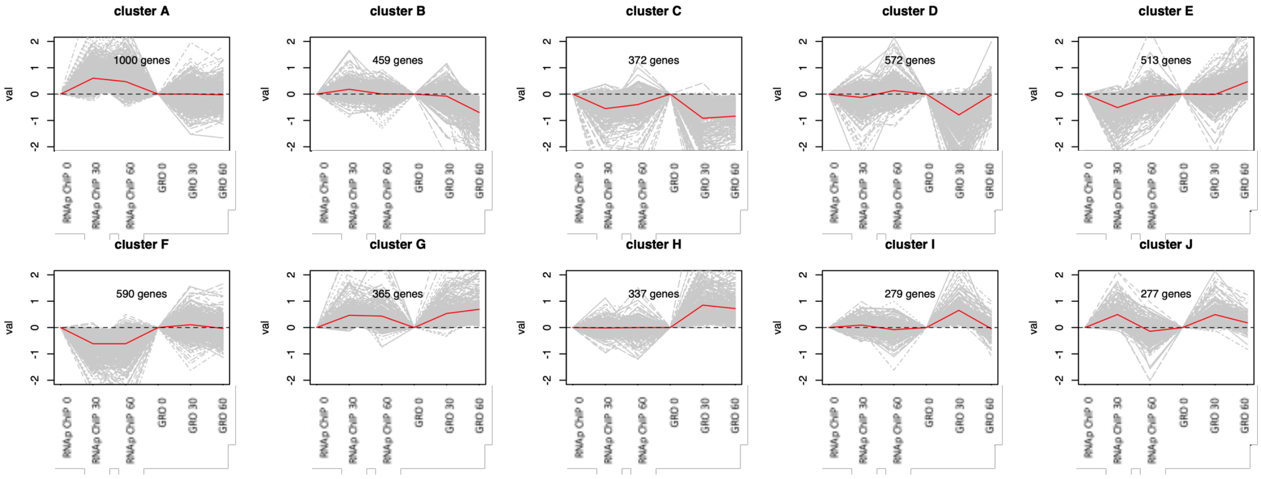

### S3 Fig

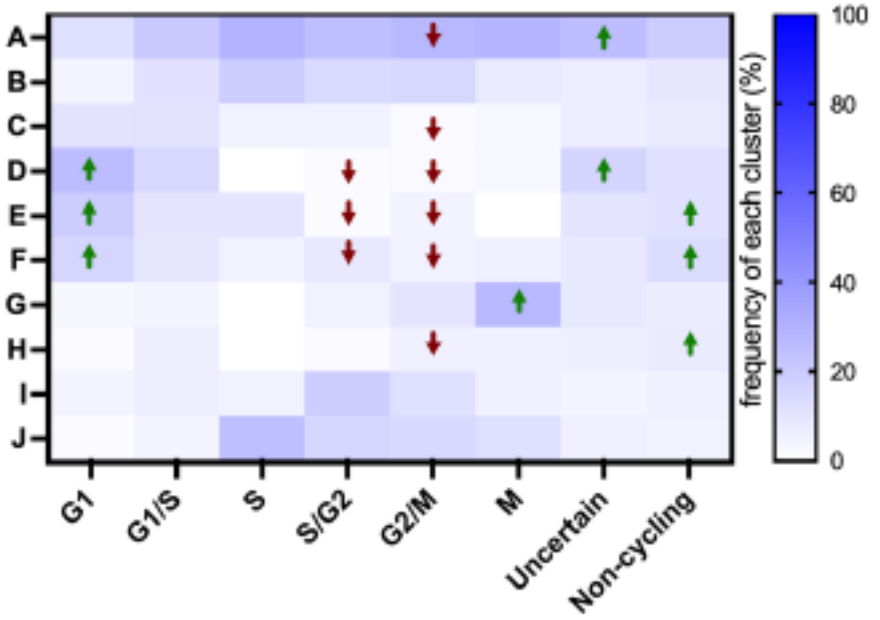

### S4 Fig

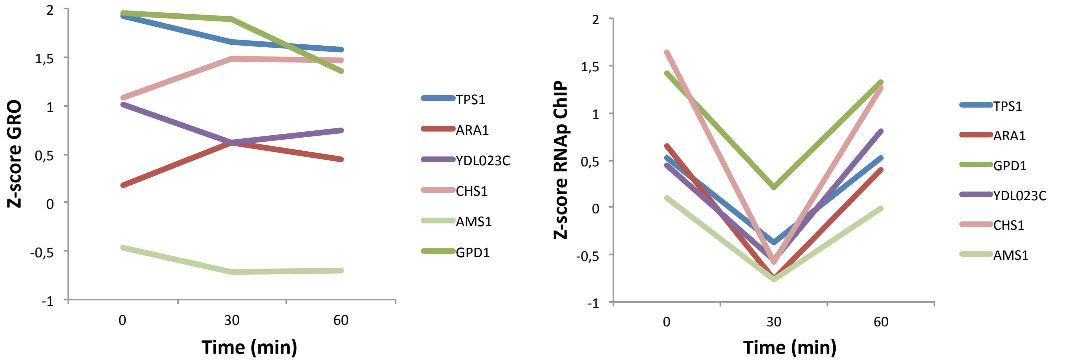

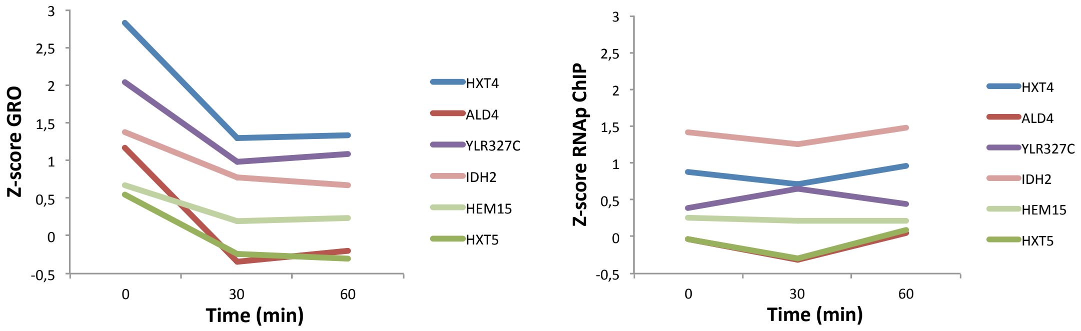

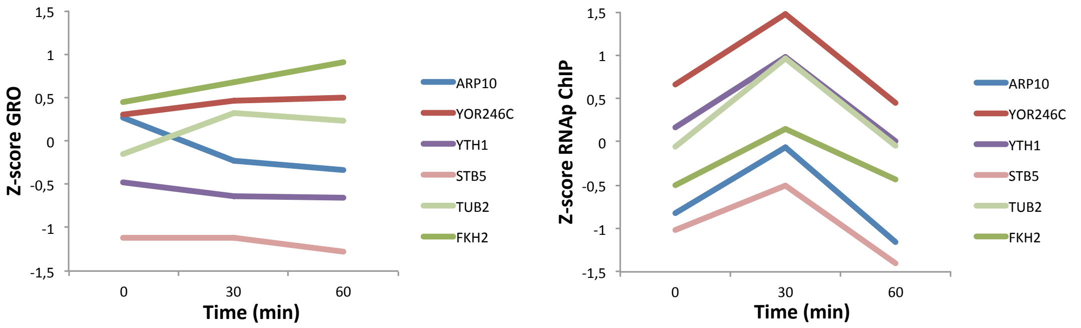

### S5 Fig

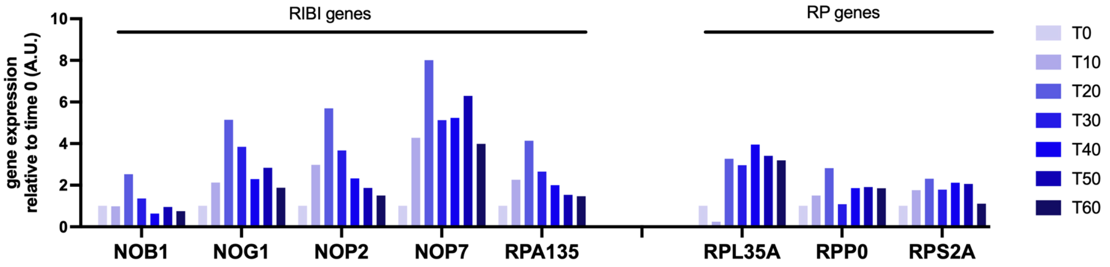
